# A local sequence signature defines a subset of heterochromatin-associated CpGs with minimal loss of methylation in healthy tissues but extensive loss in cancer

**DOI:** 10.1101/2022.08.16.504069

**Authors:** Dror Bar, Lior Fishman, Yueyuan Zheng, Irene Unterman, Devorah Schlesinger, Amir Eden, De-Chen Lin, Benjamin P. Berman

## Abstract

Global loss of DNA methylation in mammalian genomes occurs during aging and cancer, primarily in heterochromatin-associated Partially Methylated Domains (PMDs). It has previously been shown that local sequence context (100bp) has a strong influence on the rate of demethylation of individual CpG dinucleotides within PMDs. Here, we train a deep learning model to capture this sequence dependence, finding that methylation loss in healthy tissues and cancer can be predicted with high accuracy based on the 150bp surrounding a CpG. We use a published whole-genome map of the re-methylation rate of newly synthesized DNA during mitosis to show that CpGs with a “slow-loss” sequence context are efficiently re-methylated, while CpGs with a “fast-loss” sequence context are inefficiently re-methylated. Intriguingly, we find that the 10% most slow-loss CpGs lose almost no DNA methylation in healthy cell types, but lose significant DNA methylation in many cancer types. This finding suggests that loss of DNA methylation at slow-loss CpGs could underlie some cancer-specific transcriptional deregulation that has been linked to DNA hypomethylation, including the derepression of cancer antigens and transposable elements.

## Introduction

Global DNA hypomethylation was discovered as one of the epigenetic hallmarks of cancer nearly 40 years ago (*1, 2*). While its roles in cancer initiation and progression remain unclarified, it has been hypothesized to drive cancer by activating transposable elements (*3, 4*). In vitro, hypomethylation-dependent expression of TEs can also trigger a viral mimicry response and lead to a downstream innate immunity response (*5, 6*), although in vivo this may be counteracted by downregulation of innate immunity genes (*7*). Other repressive mechanisms such as histone 3 lysine 27 trimethylation can compensate for loss of DNA methylation (*8*), which may render hypomethylated tumors sensitive to epigenetic therapies such as EZH2 inhibitors (*9*).

Cancer hypomethylation occurs primarily within long regions called PMDs that are localized to the nuclear lamina and coincide with regions of late replication timing (*8, 10–12*). PMD methylation loss begins during normal aging in most tissues, and is associated with the chronological age of the individual (ZHOU). Long-term passaging in vitro has shown that methylation loss is linearly related to the number of cell divisions (*13, 14*), and the degree of hypomethylation in tumors is correlated with the number of somatic mutations (ZHOU). In non-cancer cells, all evidence to data suggests a mitotic clock-like process. In cancer, both location and degree of hypomethylation can be modulated by cancer mutations in epigenetic modifier genes such as NSD1 and DNMT3A (*15–17*), TET2 (*18*), as well as changes in cancer metabolism (*9*). PMD hypomethylation in cancer is strongly associated with hypermethylation of Polycomb-regulated CpG island promoters, which primarily occur within PMDs (*10, 12, 19*). It is unknown whether there is a mechanistic link between these two events, or whether the association is simply due to their mitotic clock properties.

Within PMDs, there is a high degree of variance of DNA methylation levels between individual CpGs. An early report showed that a great deal of this variance was due to the local (100bp) sequence context of the CpG (*20*). We showed that this was a universal feature of PMDs in both normal tissues and cancers, and that a subset of CpGs with the “solo-WCGW” context exhibited a high degree of loss across all human and mouse samples. Here, we use a deep learning approach to better characterize the sequence dependence of PMD methylation loss, and show that loss rate can be almost entirely predicted by sequence context. We show that this sequence context underlies the re-methylation efficiency of newly synthesized DNA during cell division, and that this context separates CpGs that lose methylation during normal aging from those that only lose methylation in cancer.

## Results

We first defined fast vs. slow loss CpGs within PMDs using two independent sets of human whole-genome bisulfite sequencing (WGBS) data. The “Pan-tissue WGBS dataset” included almost 400 samples from both cancer and non-cancer cell types, while the “Intra-tumor WGBS dataset” contained about 400 epithelial cells taken from several tumor samples from a human patient. These two datasets differed in cell types, genetic background of donors, WGBS protocol and sequencing coverage, and source institution. We reasoned that if machine learning models trained on each of these two datasets could converge on a single set of sequence features, those features could be considered universal.

### Pan-tissue WGBS dataset

The first dataset was a collection of WGBS datasets compiled from more than a dozen sources that included both cancer and non-cancer samples (*21*). We filtered out samples from early embryogenesis and the germline lineage, which undergo complete demethylation, and used all other cancer (n=141) and non-cancer (n=235) samples for which WGBS data was available (Supplementary Table 1). We started with all CpGs in common PMDs as defined in (*21*). In order to exclude regions of high CpG density (e.g., CpG islands), we only included CpGs with 2 or fewer other CpGs within a 150bp window (this included ~70% of all PMD CpGs). Average methylation within these 7,143,704 PMD CpGs showed a range of levels across cancer and non-cancer samples (Supplementary Figure 1A-B).

In order to identify fast-loss and slow-loss PMD CpGs, we first calculated the mean methylation level of each CpG across all 376 normal and cancer samples. Reasoning that fast-loss CpGs would have higher variance and also be more correlated to the global hypomethylation level, we calculated the covariance between each PMD CpG and the mean methylation level of all PMD CpGs across samples. Covariance had a stronger association with the known solo-WCGW than methylation mean (Supplementary Figure 1C), suggesting that covariance may be a defining feature of PMD regions. We used both the covariance and methylation mean to define 1,102,476 PMD CpGs as fast loss, and 1,756,075 CpGs as slow loss (Supplementary Figure 1C). 80% of each group were used as training data, and the other 20% were held out as test data (Figure 1A and Supplementary Figure 1D). Importantly, we also removed test set CpGs that overlapped training CpGs used for the Intra-tumor model, described below.

**Figure 1:**
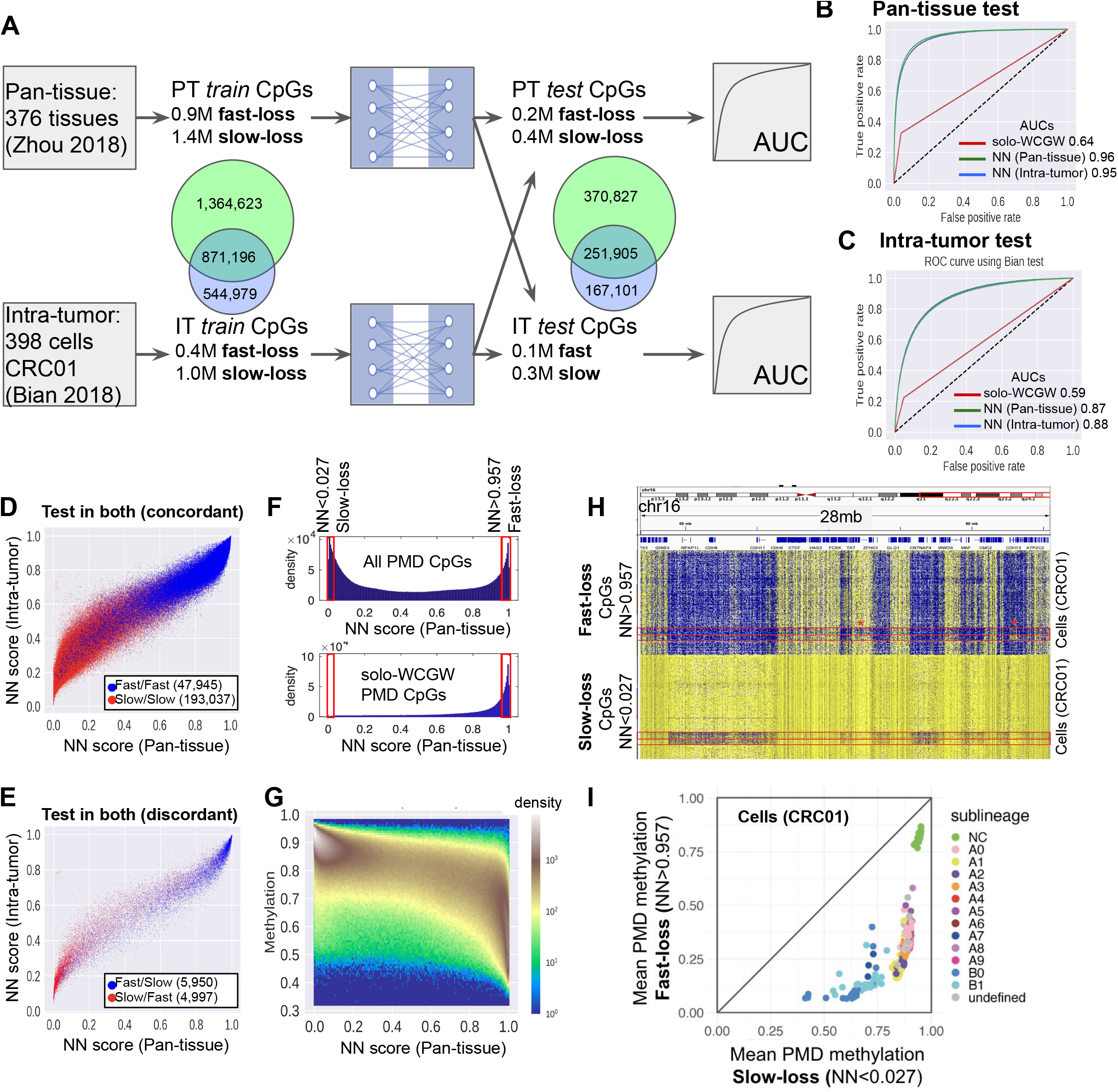
Neural network classification of fast vs. slow methylation loss CpGs. (A) A diagram of the two independent datasets used for neural network (NN) model training and evaluation. In each case, input CpGs were labeled as either “fast-loss” or “slow-loss”. For each dataset, 80% of CpGs were used for training, and 20% of CpGs were held out for testing. CpGs in either training set were excluded from both test sets. (B) NN models were evaluated using ROC analysis of the pan-tissue test set CpGs. In addition to the NN classifier, we included a binary classifier using the solo-WCGW definition from (*21*) to predict fast-loss CpGs (red line). (C) ROC analysis using the intra-tumor test set CpGs. (D) NN classification scores for 240,982 CpGs having the same fast/slow label in both test sets. (E) NN classification scores for the 10,947 CpGs in both test sets that had divergent labels (fast/slow or slow/fast). (F, top) Distribution of pan-tissue NN scores for 5,150,889 PMD CpGs (all CpGs having two or less neighboring CpGs in flanking window). Red lines are shown for the lower 10th percentile of neural network scores (0.027), and the upper 10th percentile (0.957). (F, bottom) Neural network scores for the subset of PMD CpGs labeled as solow-WCGW in (*21*). (G) Average methylation of PMD CpGs across samples in the pan-tissue dataset, as a function of NN score. (H) Genome browser plot of cells from the intra-tumor dataset (sample CRC01 from (*22*)), stratified by NN scores (fast-loss CpGs at top, slow-loss CpGs at bottom). Sublineages B0 and B1 are demarcated by red lines. (I) Cells from CRC01 sample, showing genome-wide average at fast-loss PMD CpGs vs. slow-loss. Sublineage labels were taken from (*22*), as determined by copy number alterations.

### Intra-tumor WGBS dataset

The second training dataset was from a single-cell WGBS analysis of tumor cells from a human colon cancer case (*22*). We only used one of the 10 cases sequenced (CRC01), saving the rest for validation purposes. CRC01 consisted of 389 individual flow cytometry (FACS) sorted epithelial cells from four geographically separated sections of the tumor (Supplementary Figure 1E-G). We called PMD regions for this tumor (see Methods) and identified 5,998,174 PMD CpGs. The vast majority of these (5,461,442) overlapped the PMD CpGs of the Pan-tissue dataset (Supplementary Figure 1E). In total, 71% of all PMD CpGs in either dataset were shared between the two, reinforcing the concept that most PMD regions are highly conserved across cell types, as shown previously (*10, 21*).

The original study (*22*) observed a high degree of variance of global methylation within this tumor, especially within a subclonal population of cells called “lineage B”, and we confirmed this for PMD CpGs (Supplementary Figure 1F-G). Unlike the pan-tissue WGBS set, a majority of CpGs within PMDs had methylation that was close to 0 or 1 when averaged across all cells (Supplementary Figure 1H), which is attributable to the low sequencing coverage (~0.2× per cell) and the flow cytometry selection for a pure epithelial cell population. For these reasons, the covariance metric was not used and we instead selected the CpGs close to the 0 and 1 peaks as fast-loss, and slow-loss, respectively (Supplementary Figure 1H). We verified that the “fast” CpGs were strongly associated with the solo-WCGW sequence, as expected (Supplementary Figure 1I). In total, we identified 492,208 fast CpGs and 1,342,973 slow CpGs. 80% of each group were used as training data, and the other 20% were held out as test data (Figure 1A and Supplementary Figure 1D). The test set numbers are after filtering out any CpGs that were present in the training set of the pan-tissue dataset.

The enrichment of solo-WCGW within the fast-loss CpGs in the intra-tumor dataset (25%) was not as strong as in the pan-tissue dataset (37%). This is an indication of the lower overall read coverage in the intra-tumor dataset and inability to use covariance as a selection criteria. Only about 30% of the CpGs in the two training sets overlapped (Figure 1A), compared to 71% random overlap among all PMD CpGs. This likely reflects the significant degree of noise in the intra-tumor dataset. We felt these major differences between the two training datasets would strengthen the result if the sequence models learned were highly consistent.

### Neural network classification of fast vs. slow methylation loss CpGs

The neural network model consisted of several convolutional and fully connected layers which extract information from different positions within the input sequence (Supplementary Figure 2A shows a simplified version, see Methods section for details). The input to the network is a one-hot encoded 150-bp sequence centered on a single reference CpG site. The output layer is a prediction probability of each sequence being a fast methylation loss CpG. Since all training CpGs are labeled either fast-loss or slow-loss, the probability of slow-loss is one minus the probability of fast-loss. For each dataset we trained 5 models using 5-fold cross validation and calculated the prediction of the test set using the mean of all 5 models.

To examine model performance, we used both receiver-operating characteristics (ROC) and precision-recall (PR) analysis (Figure 1B-C and Supp. Figure 3A-C). We computed ROC for each test set separately, in each case evaluating the two different models. The performance of the two models were nearly identical, suggesting that they may have converged on a nearly identical solution. We confirmed that this was the case by comparing the classification probabilities of the two models for all CpGs with the same category label in both test sets (i.e. fast/fast or slow/slow), shown in Figure 1D.

The Area Under the ROC curve (AUROC) was much higher using the pan-tissue test set (Figure 1B, AUROC=0.95) than the intra-tumor test set (Figure 1C, AUROC=0.87). This reflects the more than one order of magnitude higher sequence coverage in the pan-tissue dataset, which allowed us to use covariance to define fast vs. slow loss with significantly less noise. Out of the 251,929 CpGs in both test sets, about 4% had divergent category labels in the input (i.e. either fast/slow or slow/fast). For these divergent CpGs, both NN classifications almost always confirmed the pan-tissue dataset label, confirming this as the less noisy input dataset (Figure 1E). It is remarkable that despite this relatively high degree of noise, the intra-tumor NN model was able to converge to the same correct solution as the pan-tissue dataset. Since the two models are nearly identical, we used the pan-tissue NN model for all subsequent analyses, and recommend this as the default model for future studies.

We next looked at NN classification scores for all PMD CpGs in the pan-tissue dataset. While the NN classification scores were continuous from 0-1, the bulk of scores occurred at the two extremes (Figure 1F, top). The lowest 10th percentile (i.e. slow-loss CpGs) had scores below 0.027, while the highest 10th percentile (i.e. fast loss CpGs) had scores above 0.957 (red lines in Figure 1F). As expected, most solo-WCGW CpGs from (*21*) were classified as fast-loss (Figure 1F, bottom). However, more than half of them were not within the top 10th percentile of NN classification scores, underscoring that solo-WCGW is a relatively poor classifier of fast-loss (also shown in ROC curves Figure 1B-C). As indicated by earlier reports (*20*, *21*), there were many sequences with intermediate scores between the peaks of slow-loss and fast-loss. Methylation levels of these CpGs were also correlated to the NN score (Figure 1G). While it is possible that the intermediate signatures could be better captured by training an NN model to predict a continuous methylation level rather than the two extremes, studying the two extremes is useful for gaining an understanding of the progression of DNA methylation loss. Therefore, for all subsequent analyses, we used the top 10% of all PMD CpGs (495,640) as a set of “fast-loss CpGs”, and the bottom 10% (495,639) as “slow-loss CpGs”. These CpG positions are also available as a supplemental data file.

To visualize the methylation of fast-loss vs. slow-loss CpGs, we show a chromosome plot of the single-cell CRC01 sample, separating fast-loss CpGs (Figure 1H, top heatmap) from slow-loss CpGs (Figure 1H, bottom heatmap). At the slow-loss CpGs, the PMDs remain nearly fully methylated, except in a small subset of cells corresponding to the B0 and B1 lineages (as determined from copy number alterations in (*22*)). Plotting each of these cells as an average of all the fast-loss PMD CpGs vs. the slow-loss PMD CpGs, it is clear that most cancer cells lose methylation at fast-loss CpGs relative to the normal epithelial cells (green), but only the B0/B1 lineages lose significant methylation at the slow-loss CpGs (Figure 1I). This suggests a new paradigm where methylation loss occurs almost ubiquitously at fast-loss CpGs, but only in severe circumstances at slow-loss CpGs.

### Rate of methylation loss in normal and cancer samples revealed by sequence model

Next, we reanalyzed several independent WGBS datasets to validate the pan-tissue NN sequence model. In each case, we plotted the average of all the fast-loss PMD CpGs vs. the slow-loss PMD CpGs. We first analyzed the nine other colon cancer cases from the single-cell WGBS study (*22*) which were not used for training (Figure 2A-C, Supplementary Figure 4). We found that 3 of these 9 cases had cancer cells with significant hypomethylation of both fast-loss and slow-loss CpGs, including CRC11 and CRC13 (Figure 2A-B) and CRC04 (Supplementary Figure 4). Another 5 of 9 tumors only exhibited loss of methylation only in the fast-loss CpGs, including CRC10 (Figure 2C) and CRC02, CRC09, CRC12, and CRC15 (Supplementary Figure 4). In addition to the training sample, one other tumor had a subset of cells that became hypomethylated at slow-loss CpGs (CRC14). In all cases, the NN model classifications were validated, as fast-loss CpGs always lost more methylation than slow-loss CpGs (i.e., all cells were below the identity line). This included all non-cancer epithelial cells as well (shown as “NC” with green dots).

**Figure 2:**
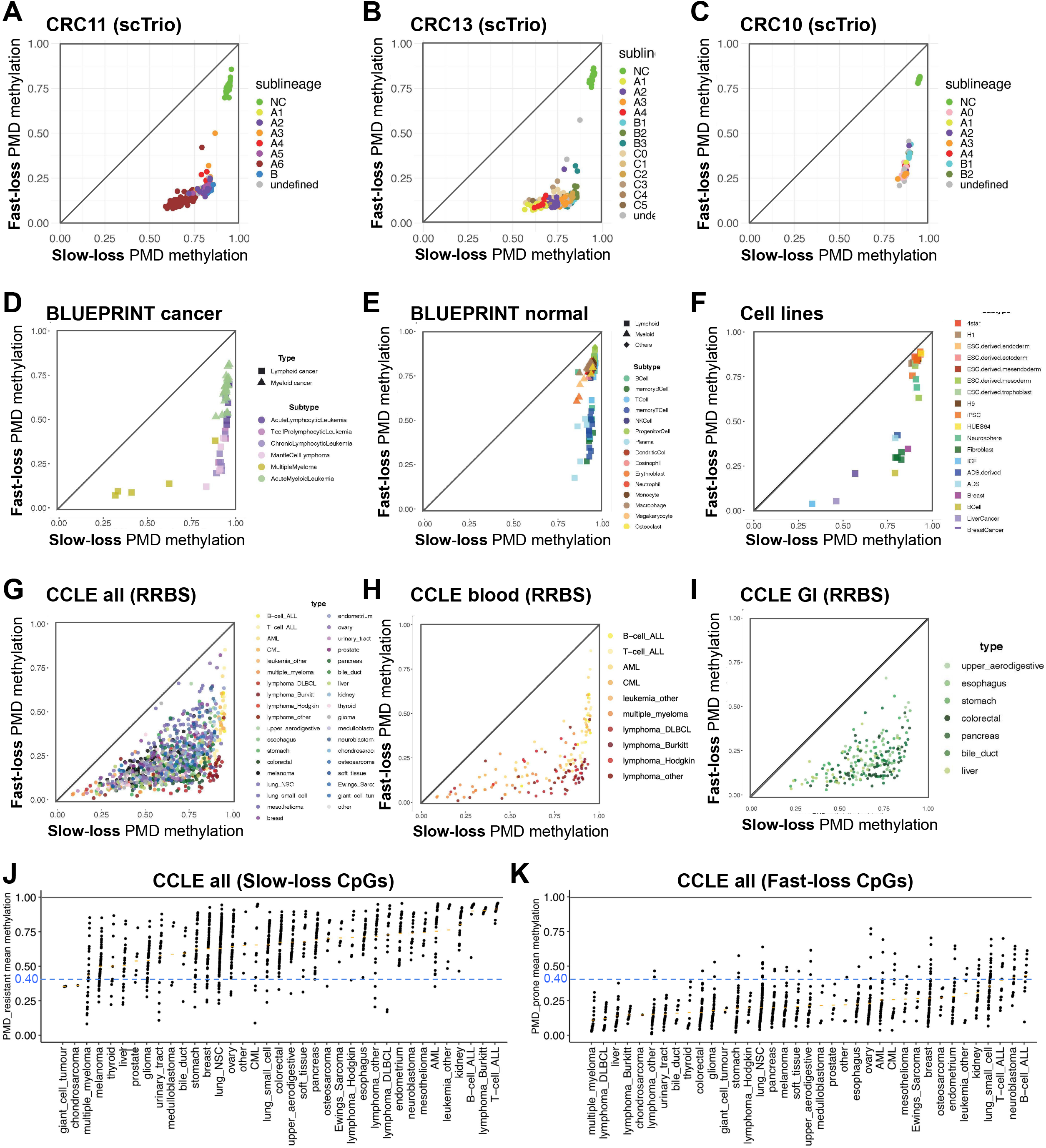
Rate of methylation loss in normal and cancer samples revealed by sequence model. (A-C) Cells from single-cell WGBS cases, showing genome-wide average at fast-loss PMD CpGs vs. slow-loss. Sublineage labels were taken from (*22*), as determined by copy number alterations. Additional cases are shown in Supplementary Figure 4. (D-F) PMD methylation plots for pure cell populations in (D-E) BLUEPRINT WGBS samples from (*23*), and (F) other WGBS samples from (*21*). (G-H) PMD methylation plots for cancer cell lines from the Cancer Cell Line Encyclopedia (CCLE) RRBS dataset (*25*). (G) shows all cancer types, (H) shows the subset of cell lines from blood cancers, and (I) shows the subset from gastrointestinal cancers. (J-K) PMD methylation for CCLE cell lines, shown separately for slow-loss CpGs (J) and fast-loss CpGs (K).

We next analyzed WGBS data from primary hematopoietic patient samples from the BLUEPRINT project (*23*). While these samples were included in the pan-tissue training dataset, they were not the majority of samples and made up very few of the samples with the most strongly hypomethylated PMDs. Most of the hematopoietic cancers had extensive loss of methylation at fast-loss CpGs but very little at slow-loss CpGs (Figure 2D). The outlier was multiple myeloma, which is known to have an exceptional degree of methylation loss (*24*).Hematopoietic cell types from healthy individuals did not have significant loss at slow-loss CpGs, even for plasma cells which are recognized to have significant loss of methylation (*21, 24*) (Figure 2E). In each case, the validity of the sequence model was verified, with fast-loss CpGs always losing more methylation than slow-loss CpGs. This was also true for cell line WGBS samples from (*21*) (Figure 2F).

We used Reduced Representation Bisulfite Sequencing (RRBS) available for the Cancer Cell Line Encyclopedia (CCLE) collection (*25*) to analyze PMD hypomethylation in a spectrum of cancer types, with none of these 928 samples were included in our training datasets. While RRBS data is biased with respect to sequence composition, we were nevertheless able to validate the NN sequence model, with fast-loss PMD CpGs losing more methylation than slow-loss PMD CpGs (i.e. below the identify line) in every single CCLE sample (Figure 2G and Supplementary Figure 6). As in the BLUEPRINT samples, dramatic hypomethylation of slow-loss CpGs occurred in multiple myeloma (Figure 2H), as well as many cancers of the digestive tract (Figure 2I). Only 11 cancer types had a median slow-loss CpG PMD methylation below 0.6, while all 37 had fast-loss PMD CpG methylation below 0.4 (Figure 2J-K). However, caution should be taken in interpreting these results as purely *in vivo* differences, given that these are cell lines that have been maintained by *in vitro* culturing.

### Relative contributions of local sequence features

We investigated the relative contributions of specific sequence features by *in silico* mutation of different sequence positions within the flanking window of test set CpGs. Specifically, we mutated between 1-3 flanking positions on either side of the reference CpG (called “flank1”, “flank1,2”, and “flank1,2,3”). We also mutated all other CpGs within the input window (Figure 3A). We found that the flank1 positions had by far the strongest influence on sequence score, with a reduction in the AUROC of 0.955-0.740=0.215 (Figure 3B). This is not surprising, given the WCGW vs. SCGS preference described in our previous work (*21*). Flanking positions had decreasing effects (flank1,2 with a further reduction of 0.047 and flank1,2,3 a further reduction of 0.021). As expected, removing all CpG neighbors also had a significant effect, with a further reduction of 0.043. An additional 0.129 of AUROC remained unaccounted for, possibly explained by positioning of the CpG relative to nucleosome positioning signals (*20, 26*).

**Figure 3:**
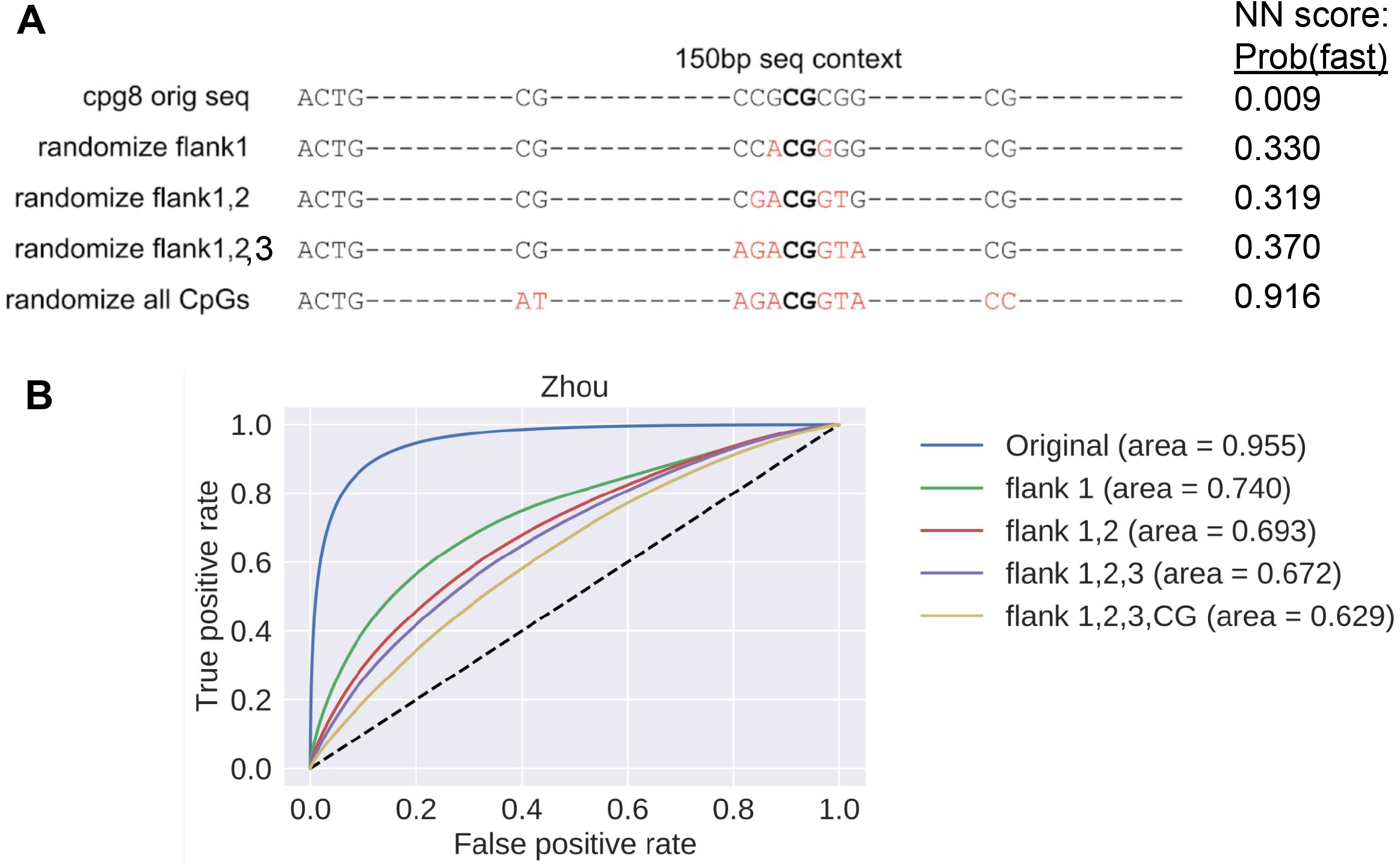
Relative contributions of local sequence features. (A) Illustration of *in silico* mutation of a test set CpG window, with the reference CpG in the center. Original nucleotides are shown in black letters, and replacements are shown in red. We show examples of each of the four mutation conditions: flank1 mutates the single base pair flanking either side of the central CpG, flank1,2 mutates the two flanking base pairs, and flank1,2,3 mutates the flanking three base pairs. The final mutation condition mutates flank1,2,3 plus all other CpG dinucleotides in the sequence. For actual in silico mutation experiments each mutation position is randomly replaced with one of the other 3 base pairs (see Methods). (B) ROC curve of pan-tissue test set CpGs classified by pan-tissue NN model, showing each successive *in silico* mutation step starting with flank1 and adding additional mutations.

### Methylation efficiency of newly synthesized DNA during mitosis underlies sequence signature

Recently, a new method to directly sequence the methylome of individual mother and daughter strands of newly replicated DNA was used to quantify methylation maintenance during cell division (*27*). Overall, the study found that DNA re-methylation mostly occurred within the first 30 minutes following replication, but was not completed until after a full 24 hours. We re-analyzed the HeLa cell re-methylation data from this study by grouping CpGs according to their NN classification scores, finding that re-methylation efficiency was perfectly correlated (Figure 4A). We also plotted the set of solo-WCGW CpGs (dotted line), which fell between the 90-95th percentile of fast-loss scores. An important limitation to this assay is that it was restricted primarily to measuring re-methylation within the most early-replicating DNA, which generally doesn’t include PMDs which are late-replicating. While we can not explicitly quantify the efficiency within PMD regions, this remains very strong evidence that re-methylation efficiency after replication is the direct cause of cumulative methylation loss within PMDs in aging and cancer, in addition to the genome wide profiling of methylation during serial passaging of cells in vitro (*13, 14*). The proposed process is illustrated in Figure 4B.

**Figure 4:**
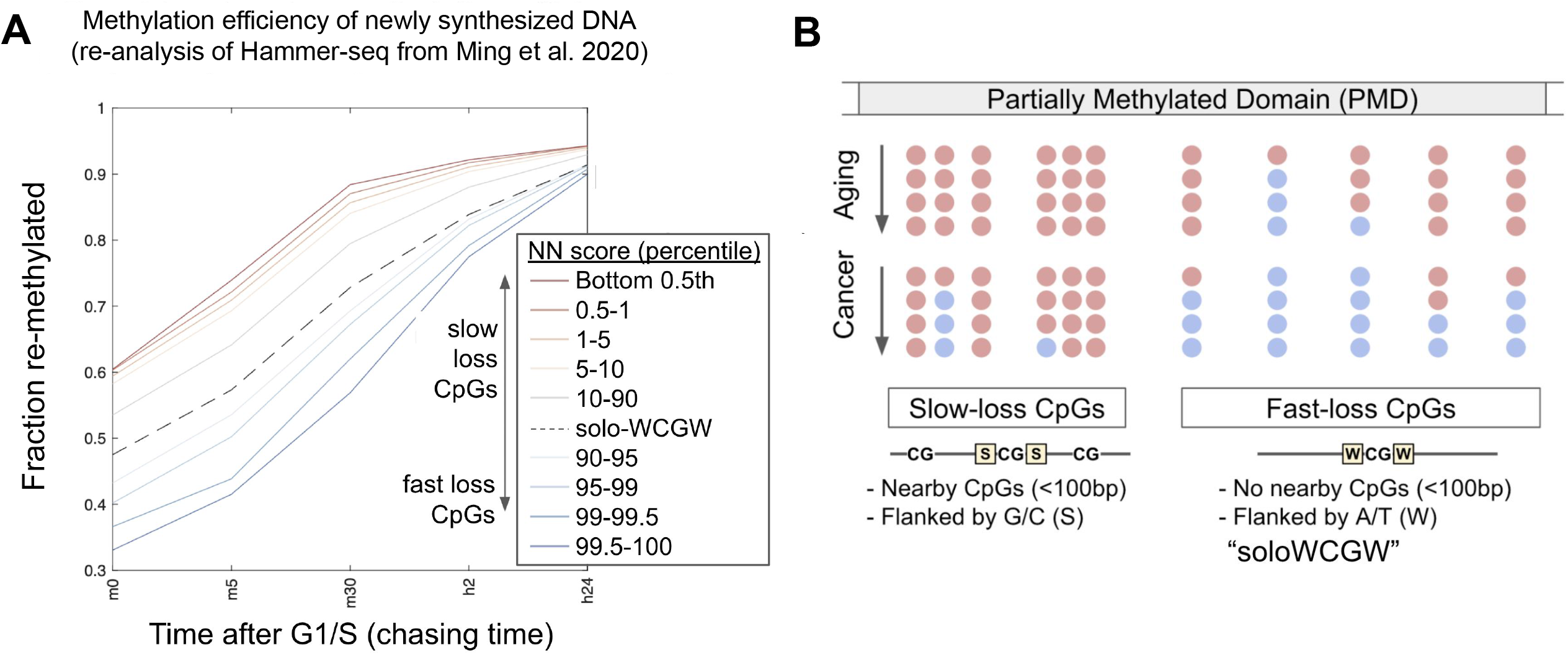
Methylation efficiency of newly synthesized DNA during mitosis underlies sequence signature. (A) Remethylation of newly synthesized DNA following replication in HeLa cells was monitored using Hammer-seq data downloaded and plotted as in (*27*). The data was re-analyzed to calculate the average remethylation as sets of CpGs defined by neural network sequence score, at the NN score percentile bins indicated. A separate dotted line shows the set of CpGs defined by the solo-WCGW sequence criteria from (*21*). (B) A model for PMD methylation loss during aging and cancer, based on local sequence features defined by the neural network score, as well as methylation state of neighboring CpGs.

## Discussion

Here, we have furthered our understanding of the forces that shape epigenetic degeneration in human aging and cancer. By using a convolutional neural network paradigm, we were able to show that slow-loss CpGs and fast-loss CpGs are nearly completely predicted based on a 150bp flanking sequence, and that these signatures predict the degree of methylation loss both during aging and cancer, in all cell types investigated. By analyzing direct measurements of nascent DNA re-methylation during mitotic S-phase, we show that this process is mediated via re-methylation efficiency of this sequence signature. These results are consistent with recent work showing that PMD methylation loss accumulates linearly with cell divisions in various human cell types (*14*).

One major question is how these sequence signatures are translated into mitotic re-methylation efficiency. We speculated previously that the CpG density component of these signatures is likely due to the processivity of the DNMT1 protein complex (*21*). If this is true, then the methylation efficiency of a daughter strand CpG may be influenced not only by the base paired CpG, but also by other neighboring CpGs, a concept recently termed “neighbor-guided correction” (*28*). There is also a strong influence of the 2-3 nucleotides flanking the CpG, and this may be due to sequence-specific DNA binding preferences of the DNMT1 protein machinery, although they do not appear to be explained by the in vitro binding preferences of DNMT1 in isolation, and could be due to the preferences of other factors such as UHRF1 (*29*).

The most important new finding here is that there are at least half a million CpGs within PMDs that have almost no methylation loss in healthy human cells, but can have extensive loss in cancer. Regions that contain these “slow-loss” CpGs thus may be potential candidates for regulatory elements that become dysregulated in strongly hypomethylated cancers. A number of transcription factors are repressed by DNA methylation (*30, 31*), and these may be unlocked as transcription activators for binding sites that include slow-loss CpGs. Such events may underlie aberrant expression of transposable elements (TEs) or cancer testis antigens (CTAs). While both of these transcript classes are broadly upregulated in cancer as well as in normal cells upon treatment with chemical demethylating agents, a direct regulatory link with global methylation loss *in vivo* has not been established. Interestingly, interferon genes that respond to endogenous TE expression are downregulated in strongly hypomethylated cancers (*7*), perhaps as a survival mechanism to counteract immune responses. Interestingly, a recent study showed that aberrant expression of CTA genes in breast cancer could be attributed to a replication-linked methylation loss clock (*32*), likely corresponding to the signature described here. Because this process affects lamina-associated PMDs much more severely than other domains, it will be important to establish that any regulatory consequences are specific to loci within these PMDs.

Another interesting question about methylation loss at these slow-loss CpGs is how they may influence regulation of CpG island promoters. Due to high CpG density, most CpGs within CGI promoters are inherently slow-loss. While most CGI promoters are fully unmethylated in normal cells, a subset with intermediate CpG density become highly methylated (*33*). Furthermore, the intermediate density “shores” of CpG islands are much more cell-type specific (*34*). It will be interesting to investigate whether these regions are susceptible to hypomethylation-associated activation specifically in cancer, since many of these regions contain slow-loss CpGs. Furthermore, a subset of CGI promoters regulated by the Polycomb Repressive Complex 2 become hypermethylated in cancer, and it will be interesting to investigate whether this process can be counteracted by hypomethylation at slow-loss CpGs.

Since mitotic methylation loss is a type of molecular “clock”, it is important to put our work in the context of other methylation clocks. Our PMD-based hypomethylation signature was recently confirmed as a mitotic clock in vivo in B-cell development and cancer (*24*) and in cell divisions in vitro in several human cell types (*14*). Each of these studies defines their own minimal versions of a hypomethylation clock, “epiCMIT” and “RepliTali”, respectively. Our goal here is different - to better understand the mechanisms rather than design a minimal clinical signature. Because DNA hypomethylation is associated with chronological age (*21*), it is important also to consider methylation clocks discovered directly from age association, such as the Horvath pan-tissue aging clock (*35*) and derivatives. We showed previously that these clocks contain very few PMD CpGs associated with our mitotic signature, likely due to their reliance on the Infinium methylation array platform which has a strong underrepresentation of these sequences (*21*), and because they are only weakly associated with chronological aging.

## Code availability

Source code for neural network analysis at https://github.com/methylgrammarlab/pmd_hypometh_classifier. Source code for calculating the mean methylation of fast and slow-loss CpGs of CCLE and Pan-tissue WGBS dataset is in https://github.com/yuanzi2/FastSlowScatterPlot. Scripts to process hammerseq data are available at https://github.com/methylgrammarlab/NN_hypomethylation_hammerseq.

## Data availability

Processed data files for the analyses described here are available at Zenodo accession 10.5281/zenodo.6592632.

## Acknowledgments

We thank Yontanan Berg, Amit Levin, Dolev Revivo for useful discussions regarding PMD hypomethylation analysis. We thank Tiago Silva for help with R/Bioconductor programming. Ben Berman received startup support from the Hebrew University, the Kamea B program of the Israel Ministry of Aliyah and Immigrant Integration, and the Israel Cancer Research Fund Project Grant (845755). Irene Unterman received support from the Data Sciences Initiative from the Council for Higher Education in Israel (MALAG) and the Hebrew University Center for Interdisciplinary Data Science Research (CIDR).

## Author Contributions

DB and LF collected and processed training and test data and implemented neural network classifiers. YZ collected validation data and performed validation analysis. IU, DS, and AE performed additional validation analysis. BPB conceived the project. BPB and DCL supervised analysis. BPB, DB, LF, and DCL wrote the manuscript. The first three authors (DB, LF, YZ) are co-lead authors with an equal contribution to the work, and they have the right to list their names first in their CVs. The last two authors (DL,BPB) are co-senior authors who jointly supervised the work, and they have the right to list their names last in their CV.

**Supplementary Figure 1:**
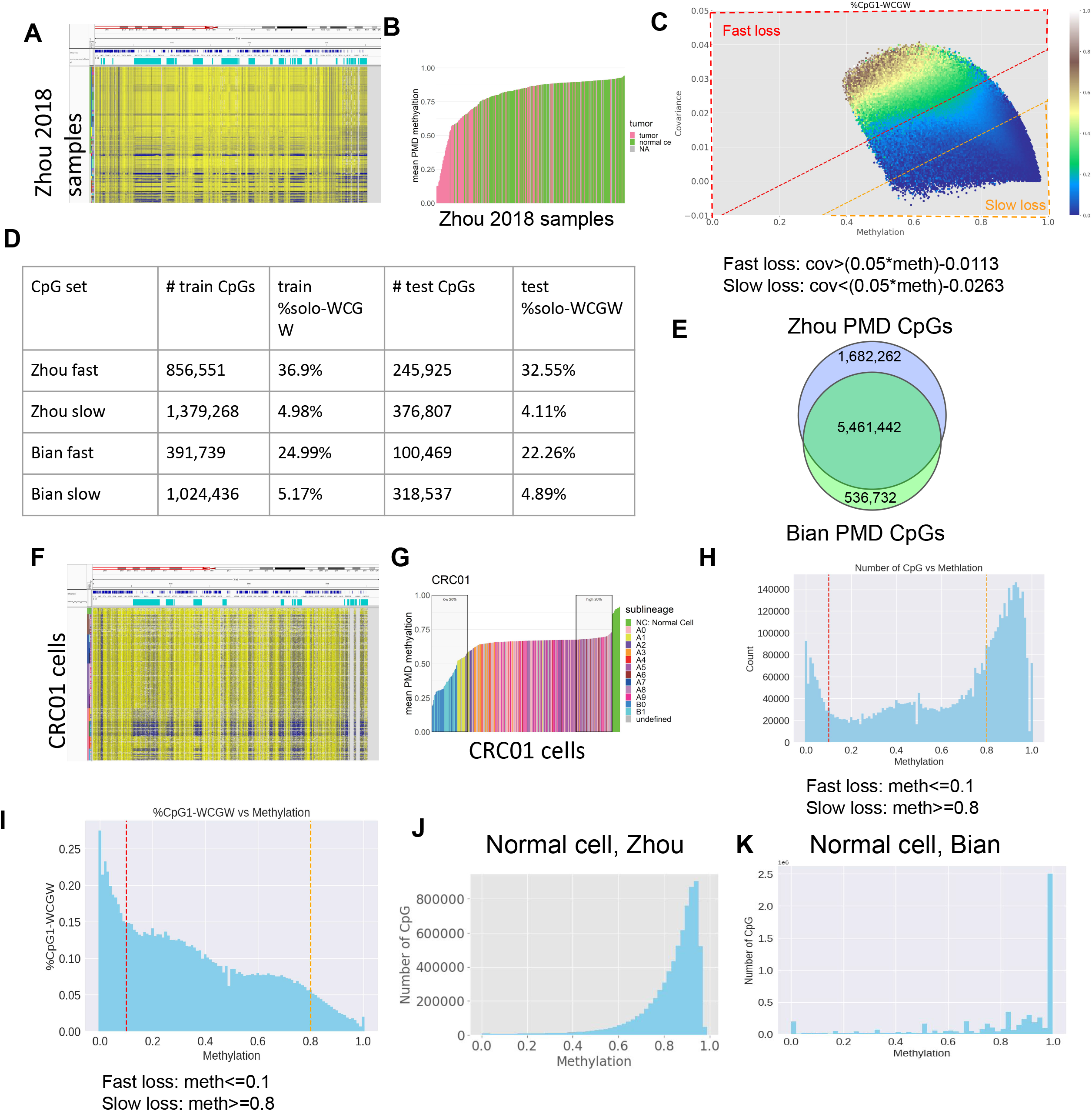
Creating training and test data from two WGBS datasets. (A) Methylation values for all samples in the pan-tissue dataset showing chromosome arm 16p, with Partially Methylated Domains (PMD) locations taken from (*21*), and yellow indicating high methylation. (B) Average methylation levels within PMDs for each sample, colored by cancer versus non-cancer. (C) A heatmap of PMD CpGs from the pan-tissue dataset (only those with 2 or fewer neighboring CpGs within 150bp), plotted on an x-axis representing the average methylation across all samples, and a y-axis representing the covariance between the CpG methylation values across samples and the global PMD methylation averages across samples. The color of the heatmap represents the percentage of CpGs near the x,y coordinate that match the solo-WCGW definition from (*21*). Red and orange boxes show the CpGs picked as fast-loss (red) and slow-loss (orange) for the pan-tissue training and test sets. The cutoff for the fast-loss group is covariance > ((0.05 * methylation)-0.0113), and the cutoff for the slow-loss group is covariance < ((0.05*methylation)−0.0263). (D) Summary table of CpG counts for pan-tissue (Zhou) and intra-tumor (Bian) training and test sets. (E) Overlap of input PMD CpGs (before selection) between pan-tissue (Zhou) and intra-tumor (Bian) datasets. (F) Chromosome 16p plot of all cells from the intra-tumor dataset (colon cancer case CRC01 from (*22*)), with yellow indicating high methylation. (G) Average methylation levels within PMDs for each cell of CRC01, colored by genetic sub-lineage. (H) Histogram of CpG counts in CRC01 based on average methylation across all cells. Red and orange lines show the cutoffs for CpGs picked as fast-loss (red, methylation<0.1) and slow-loss (orange, methylation>0.8) for the intra-tumor training and test sets. (I) Plot of the percentage of CpGs matching the solo-WCGW definition from (*21*), for bins of CpGs having the same average methylation across CRC01 cells. (J) Histogram of average methylation level of PMD CpGs only within non-cancer samples in the pan-tissue dataset. (K) Histogram of average methylation level of PMD CpGs only within non-cancer epithelial cells in the intra-tumor CRC01 dataset.

**Supplementary Figure 2:**
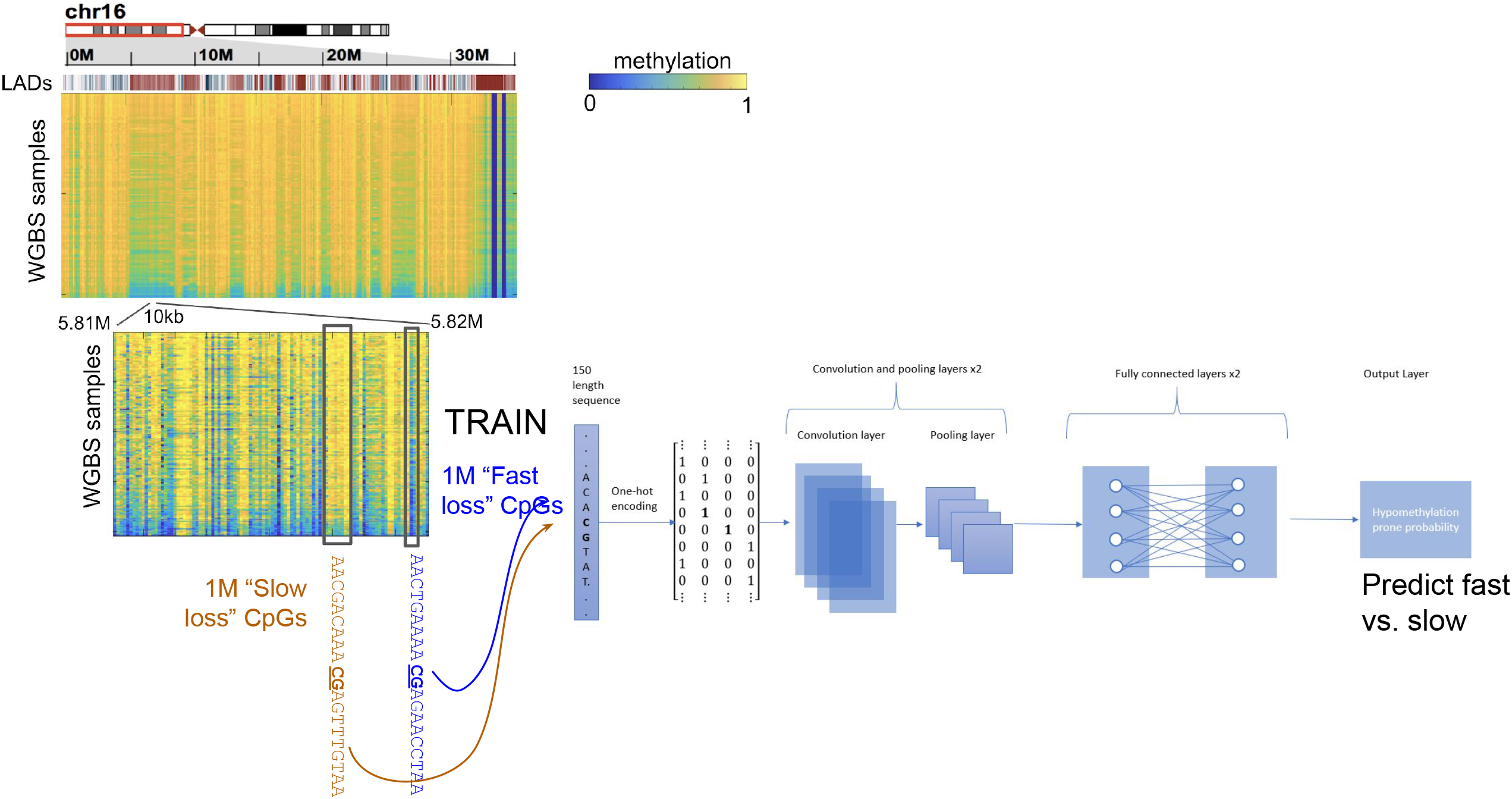
Schematic of neural network training. (A) The input for each training set is the subset of CpGs within PMDs that have 2 or less neighboring CpGs within a 150bp window. Based on average methylation and covariance across samples/cells, fast-loss and slow-loss CpGs are selected. 80% of these CpGs are used for training, and 20% are held out as a test set for model evaluation. For each training set CpG, the 150bp window centered on the CpG is one-hot encoded as the input layer for a convolutional neural network. The output layer is the fast-loss or slow-loss label, and the network predicts whether it is a fast-loss CpG (probability of 1 means complete confidence that it is fast-loss CpG, and a probability of 0 means complete confidence that it is a slow-loss CpG.) For each dataset we trained 5 models using 5-fold cross validation and calculated the prediction of the test set using the mean of all 5 models. Further details about network architecture are available in the Methods section.

**Supplementary Figure 3:**
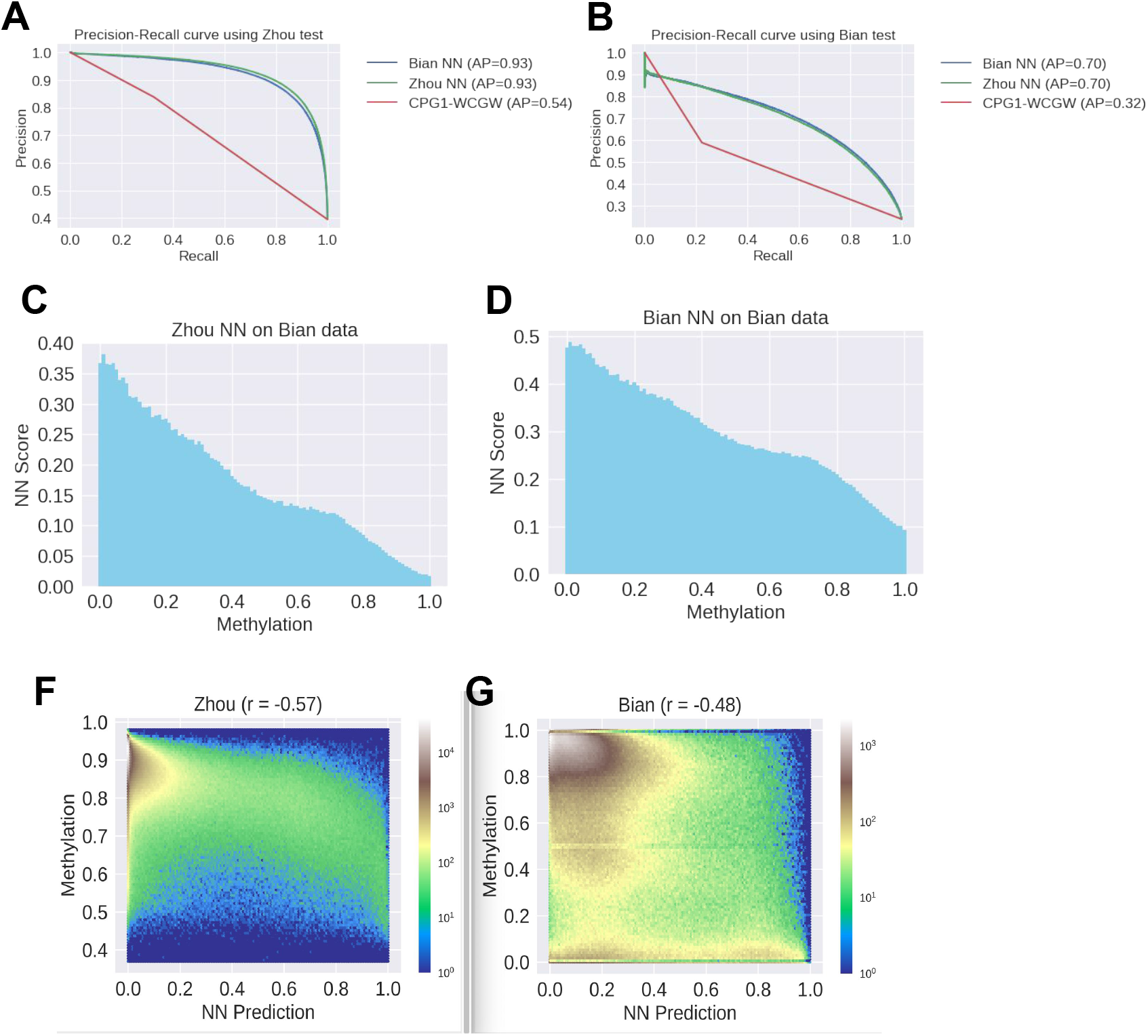
Neural network evaluation features. (A-B) Precision-recall plots are shown based on the same data as ROC plots shown in Figure 1B-C. Panel (A) corresponds to the pan-tissue test set (Zhou) evaluated on both models, and (B) corresponds to the intra-tumor test set (Bian) evaluated on both models. (C-D) NN scores for pan-tissue model (C) and the intra-tumor model (D), for bins of CpGs having the same average methylation PMD methylation averages across cells in the intra-tumor CRC01 dataset. (F) Average methylation for PMD CpGs in the pan-tissue dataset, based on NN scores from the model trained with the intra-tumor training data. (G) Average methylation for PMD CpGs in the intra-tumor dataset, based on NN scores from the model trained with the intra-tumor training data.

**Supplementary Figure 4:**
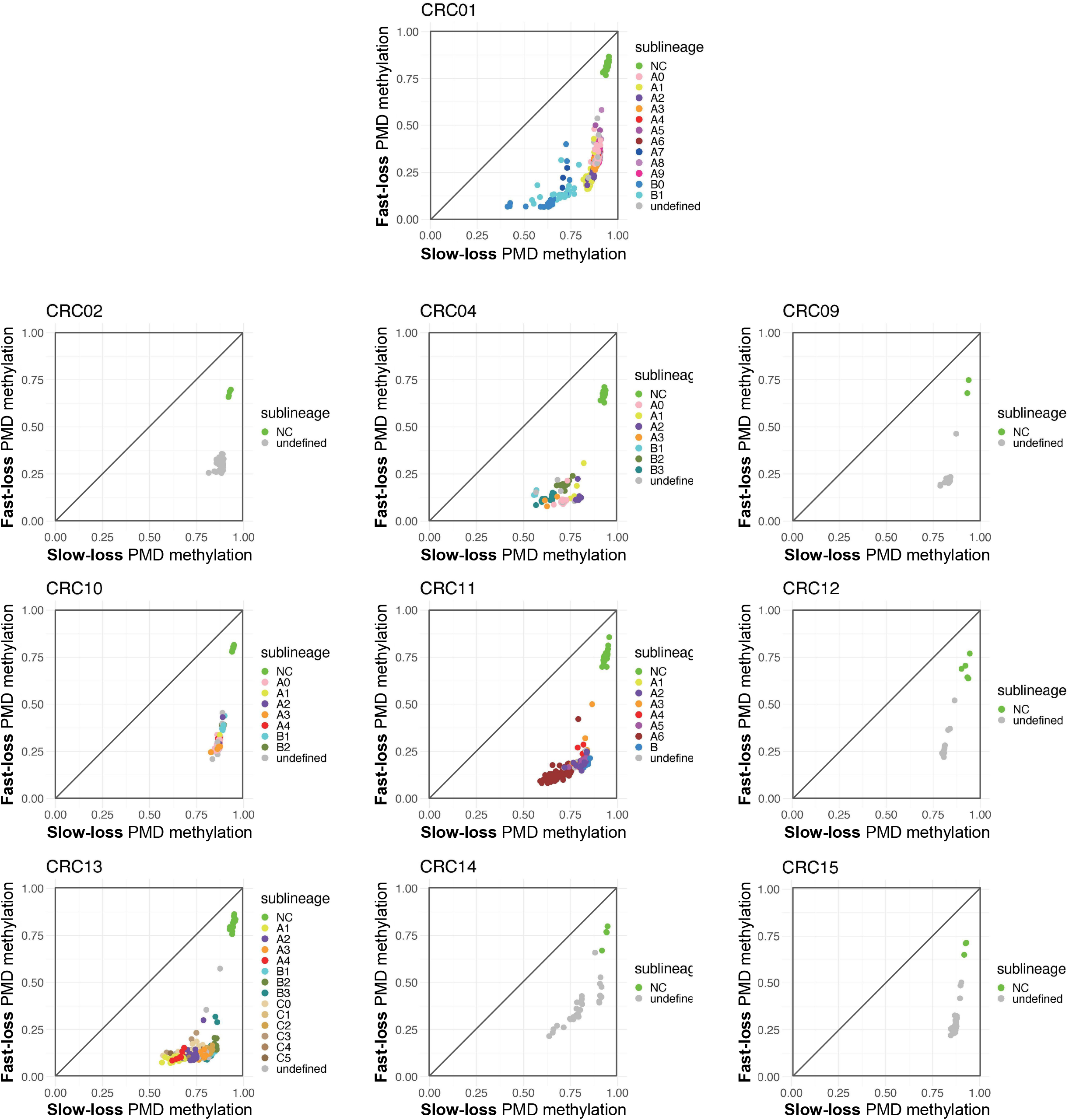
Rate of methylation loss in single-cell colon cancer data revealed by sequence model. Cells from each single-cell WGBS case from (*22*), showing genome-wide average at fast-loss PMD CpGs vs. slow-loss. Sublineage labels were taken from (*22*), as determined by copy number alterations. Several cases are duplicated from Figure 1I and Figure 2A-C.

**Supplementary Figure 5:**
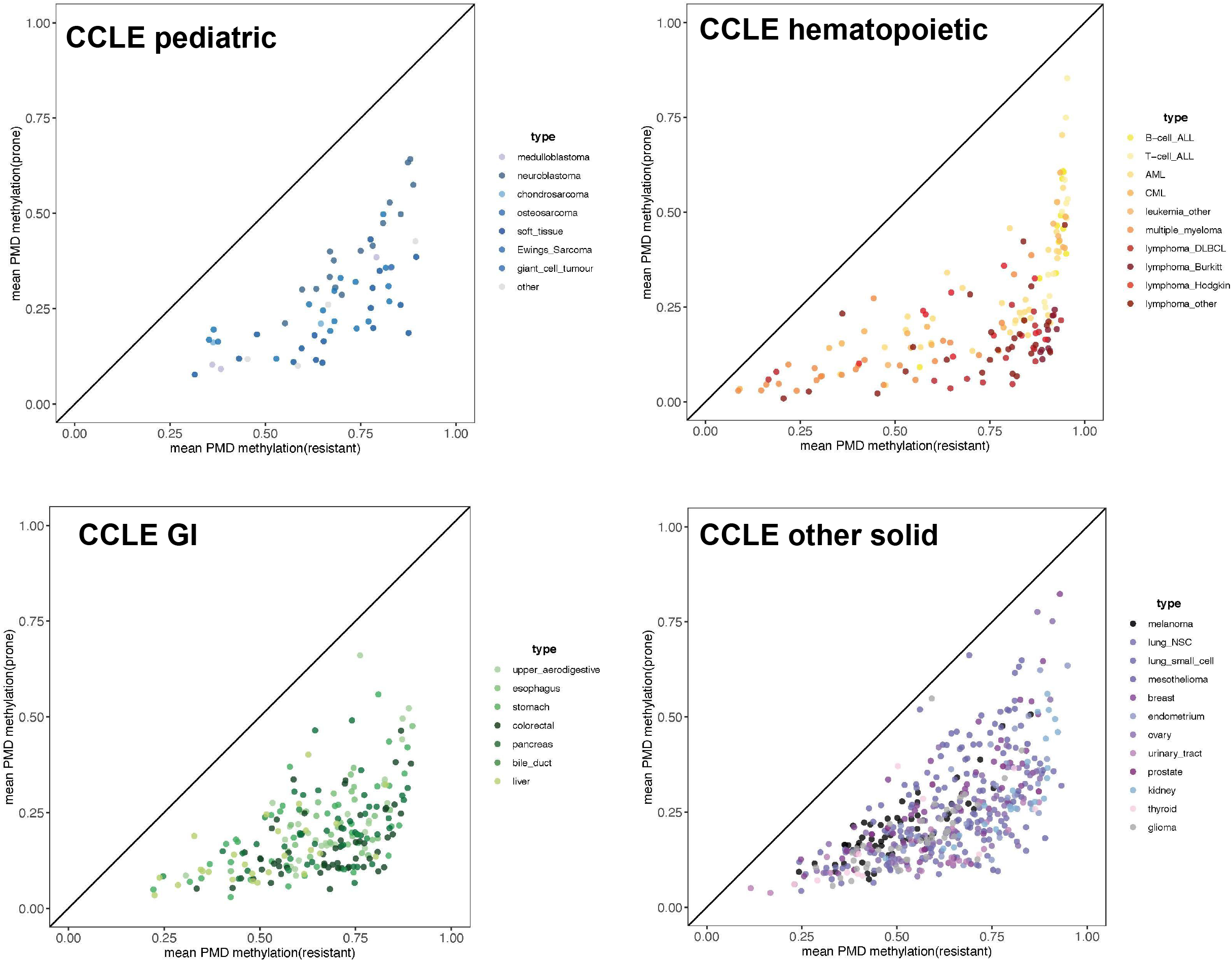
Rate of methylation loss in cancer cell lines revealed by sequence model. PMD methylation plots for cancer cell lines from the Cancer Cell Line Encyclopedia (CCLE) RRBS dataset (*25*), showing genome-wide average at fast-loss PMD CpGs vs. slow-loss. Each plot shows a different set of cell lines based on cancer type, with each dot being an individual cell line.

**Supplementary Table 1: WGBS samples from training datasets.**

**Supplementary Table 2: PMD methylation levels for WGBS and RRBS datasets.**

## Methods

### Pan-tissue training dataset

The Zhou et al. dataset includes 376 samples. 26 samples described in the original paper were not available as bed files, and they are omitted (Supplemental Table 1). This data is available from (*21*) at https://zwdzwd.github.io/pmd.

### Intra-tumor dataset

The Bian et al. dataset includes single cell Trio sequencing - Methylation (WGBS), Expression (RNA-seq) and Copy number (WGBS). The data was collected from 10 colorectal cancer (CRC) patients with between 23 to 289 cells per patient. The samples came from different tissues – the primary tumor site, lymph node metastasis, omental metastasis, liver metastasis, post treatment liver metastasis and normal tissue. The Bian data set was obtained from (*22*) using the GEO project ID GSE97693.

### Quality filtering of intra-tumor dataset

*S*ome cells in scWGBS perform poorly because of low DNA quantity or isolations containing more than one cell. We filtered out cell samples whose coverage differed by more than 2 s.d. from the mean coverage of all chromosomes across patients (mean coverage was 0.2677 and standard deviation 0.1181). Then to filter out individual regions, we filtered out CpG sites that had more than 4 reads. Some of these might represent good data do simply to cancer genome amplifications, but we chose this filter to use only high quality data. This left us with 90.3% of all data remaining. Lastly, we filtered out blacklisted CpGs (*36*). While only CRC01 was used for model building, we performed the same filtering of all ten samples from (*22*) to investigate hypomethylation dynamics (Supplementary Figure 4).

### Definition of PMD regions

All analysis was done on CpGs within PMD regions. For the pan-tissue dataset, we used common PMD regions defined in (*21*) and available at https://zwdzwd.github.io/pmd. For the intra-tumor dataset, we used only cancer cells, due to the small number of normal cell samples for some cases. We identified PMDs by covariance, which is similar to how (*21*) identified PMDs using variance. The covariance was calculated in the following manner: we split each chromosome to windows of 5000 CpG sites, for each site in the window the covariance was defined as the average covariance of that site with all other sites in the window that share at least 10 valid samples. We removed sites whose covariance was based on less than 750 pairs. In order to avoid covariance due to small technical changes, we only used the 20% of cells with the highest methylation and the 20% of cells with the lowest methylation.

### Selecting CpGs for model training and evaluation

We excluded CpG sites whose coverage was in the top or bottom 5% to remove extreme outliers. Next, we excluded sites that were unmethylated in the normal cells meaning an average methylation value lower than 0.5, this value was chosen by looking at the distribution of the methylation within PMDs (Supplementary Figure 1H-I). For the Zhou dataset this was calculated by the samples representing non-cancer tissues, and for the Bian dataset this was calculated using the average of normal cells across all the patients to increase the coverage.

For both datasets, we selected only CpGs with 3 or fewer CpGs within the 150bp window centered on and including the reference CpG (we refer to these as the CpG1, CpG2, or CpG3). The majority of common PMD CpGs (67.5%) were in this group, and those with greater than two neighboring CpGs rarely become hypomethylated (*21*). To verify that in our own datasets, we calculated the percentage of fast loss CpGs with greater than 2 neighboring CpGs, which was 6.35% and 14.15% in the Zhou and Bian datasets, respectively. In testing our NN models, we found that these sequences were not effectively learned due to their poor representation in the data, and that more sophisticated training data balancing would be required. We therefore focused on the CpGs with 2 or less neighboring CpGs.

### Criteria for fast and slow CpGs

For the pan-tissue dataset, we calculated both mean methylation across all samples, as well as covariance between that CpG and the mean methylation of all filtered PMD CpGs. Based on the enrichment for solo-WCGW (Supplementary Figure 1C), we chose fast loss CpGs as those with Cov>((0.05*meth)-0.0113) and slow loss CpGs as those with Cov<((0.05*meth)-0.0263). These are the lines drawn in Supplementary Figure 1C. All fast or slow CpGs in a PMD matching the above criteria were added to either the train or the test, with the train data including CpGs from the 7th to the 81st PMD in every chromosome and the test data included the rest.

For the intra-tumor dataset, we chose fast loss CpGs as those with methylation mean less than 0.1, and we chose slow loss CpGs as those with methylation mean greater than 0.8. These are the lines drawn in Supplementary Figure 1F. All valid CpGs in a PMD were added to either the train or the test, with the train data including CpGs from the 8th to the 82nd PMD in every chromosome and the test data included the rest.

After train and test sets were defined for both models, any CpGs included in both a training set and a test set were included from both test sets, so that test sets would be completely independent of training. The numbers of CpGs are given in Figure 1A and Supplementary Figure 1J. Of the 2,780,798 CpGs selected as fast or slow loss training examples in either model, 871,196 (31%) were shared between the two (Figure 1A). These were included as training for both models. Of the 789,833 CpGs that were selected for the test set, 251,905 (31%) were shared between the two, and these were used for evaluation of both models.

### Construction of neural network

Samples for the network were 150bp long strands with a central CpG. All sequences were based on hg19, and are encoded using one-hot encoding. We used each strand of the CpG as two independent inputs (they are exact reverse complements of each other). All figures appearing in this paper refer to the number of unstranded CpGs, but the number of input records for NN training was always doubled.

The deep neural network (DNN) architecture is based on a convolution neural network (CNN) architecture used by DeepRiPe (*37*), with adaptations to the data and to increase the network’s accuracy. The input is a sequence of 150bp long strands with a centralized CpG site (positions 74,75). The input sequence has two convolution layers followed by a rectified linear unit (Relu), an L2 regularization penalty, a max pool layer and a dropout layer with probability of 0.25. The first convolution layer uses 90 filters with kernel size of 3 and 100 filters and kernel size of 5 for the second one. The pool size after both layers is 2. The second convolution layer is connected to two fully connected layers, both with a Relu and L1 regularization penalty. The first network contains 500 hidden units and the second one 250 hiddens units. Between the two hidden layers we have another dropout layer with probability of 0.5 to prevent memorizing and overfitting for the training set. The output layer contains a sigmoid to predict the probability of a sequence to be fast loss (1) or slow loss (0).

We ran the network with a batch size of 128 with 20 epochs with an early stop option based on the validation loss to prevent overfitting during the training, and also using an Adam optimizer (*38*). An imbalanced loss penalty was used to handle the inherent imbalance in the train and test datasets labels. For both datasets the training data was split again to train and validation, based on 80% - 20% using stratified-5-fold cross validation.

### Producing scores for all hg19 CpGs using NN

As a resource for future work, we ran each CpG in the hg19 genome as an input to each neural network model, to produce a fast-loss probability score. These scores are available within Zenodo accession 10.5281/zenodo.6592632 in the file “zhou-bian.allCGs.1based.hg19.tsv.gz”, which uses 1-based coordinates and includes additional information columns “cpgi” and “blacklist”. “cpgi” has a value of 1 if it is covered by a CpG Island under the Irizarry definition (*39*). For convenience, we provide a version of this same data for the pan-tissue neural network model (the preferred model), in standard 0-based bedgraph format, in a file called “multitissue-nn-scores.allCGs.0based.hg19.bedgraph.gz”.

### Fast vs. slow methylation plots

For all figures showing mean values of “slow-loss” and “fast-loss” CpGs (Figures 1F, 1H, 1I, 2A-K), we used CpGs from PMDs of the pan-tissue dataset 3 or fewer total CpGs within the 150bp window centered on the reference CpG (the same subset of CpGs used for model training and evaluation). We then took the 10% most fast-loss CpGs based on the pan-tissue neural network model (495,640 CpGs with score>0.957), and the 10% most slow-loss CpGs (495,639 CpGs with score<0.027), as shown in Figures 1F,1H. To calculate averages for each sample, we took the average methylation level within each of these CpG sets as the average “fast-loss” and average “slow-loss” methylation, respectively. For convenience, we have provided this as an hg19 bedgraph file within Zenodo accession 10.5281/zenodo.6592632 as the file “zhou_NN_scores_PMD_top_bottom_10ptile.0based.hg19.bed.gz”.

### BLUEPRINT WGBS processing

139 blood samples and 50 blood malignancies were obtained from the BLUEPRINT data portal (ftp://ftp.ebi.ac.uk/pub/databases/blueprint/). For each sample, we downloaded the coverage and methylation bedgraph files and reserved those CpGs with a coverage >=10 reads. All files were based on hg19.

### CCLE/DepMap RRBS processing

Fastq RRBS files were downloaded for all CCLE cell lines from the DepMap website (SRA accession number PRJNA523380). We realigned and processed exactly as in https://dx.doi.org/10.1038%2Fs41586-019-1186-3. Specifically, short reads from the RRBS data were aligned using Bismark v0.23.1 for 928 cell lines. CpG methylation was estimated using the read.bismark tool in the R MethylKit package with parameters mincov = 5 and minqual = 20. Genome version hg38 was used.

### Hammerseq processing

Methylation “events” files were downloaded from the GEO series GSE131098 (*27*). We used files corresponding to the 5 time points for the wildtype HeLa-S3 sample: GSM3763438_WT-c0.events.txt.gz, GSM3763439_WT-c5m.events.txt.gz, GSM3763442_WT-c30m.events.txt.gz, GSM3763443_WT-c2h.events.txt.gz, GSM3763446_WT-c24h.events.txt.gz. These were the same 5 timepoints used in the main figures of the source publication (*27*). From these events files, we calculated a percent methylation for each genomic position represented (after removing ENCODE blacklist regions). This was done using a script “eventsToBedgraph-clean.pl” (available from https://github.com/methylgrammarlab/NN_hypomethylation_hammerseq), which exported for each coordinate a count of “MM” events (methylated in original strand and newly synthesized daughter strand), and a count of “MM+MU” events (methylated in original strand and either methylated or unmethylated in the daughter strand). This was merged with neural network scores from the file “multitissue-nn-scores.allCGs.0based.hg19.bedgraph.gz”, and Figure 4 was produced using a custom MATLAB script “READMEclean.m” (available from https://github.com/methylgrammarlab/NN_hypomethylation_hammerseq). This script extracted the subset of coordinates with neural network scores in the following bins: 0-0.5 percentile (0<=NN<=0.00076), 0.5-1 percentile (0.00076<=NN<=0.00176), 1-5 percentile (0.00176<=NN<=0.01183), 5-10 percentile (0.01183=NN<=0.02746), 10-90 percentile (0.02746<=NN<=0.95698), 90-95 percentile (0.95698<=NN<=0.98212), 95-99 percentile (0.98212<=NN<=0.99667), 99-99.6 percentile (0.99667<=NN<=0.99817), and 99.5-100 percentile (0.99817<=NN<=1.0). For each subset, the total MM events across all CpGs was divided by the total number of MU events across all CpGs. Note that this weights CpGs unequally based on their read coverage. We also tried weighting CpGs equally, and resulting plot was visually indistinguishable.

